# Correlation Coefficient Based Automatic Optic Disc Detection from Retinal Images

**DOI:** 10.1101/2020.06.23.166793

**Authors:** S. Anand, R. Prabhadevi, D. Rini

## Abstract

In this paper an algorithm to detect the optic disc (OD) automatically is described. The proposed method is based on the circular brightness of the OD and its correlation coefficient. At first the peak intensity points are taken, a mask is generated for the given image which gives the circular bright regions of the image. To locate the OD accurately, a pattern is generated which is similar to the OD. By correlating the retinal image with the pattern generated, the maximum correlation of the pattern with the OD is obtained. On locating the coordinates of maximum correlation, the exact location of the OD is detected. The proposed algorithm has been tested with DRIVE database images and an average OD detection accuracy of 95% was obtained for healthy and pathological retinas respectively.

## I. INTRODUCTION

Image processing and edge detection in in images have variety of applications [1], [2] [3], [4], [5]. Automatic OD Detection (ODD) from retinal images is essential to detect various types of eye diseases. Detecting and counting lesions in the human retina like microaneurysms and exudates is a time consuming task for ophthalmologists and prone to human error. That is why much effort has been done to detect lesions in the human retina automatically. Finding the main components in the fundus images helps in characterizing detected lesions and in identifying false positives. The detection of the OD is the first step in the early detection any eye disease. The OD is a bright circular region present in the retina. It is placed 3 to 4 mm to the nasal side of the fovea. It represents the beginning of the optic nerve and entry for the main blood vessels that source the retina. The brightness of the OD is due to the convergence of the blood vessels.

**FIG:1.**
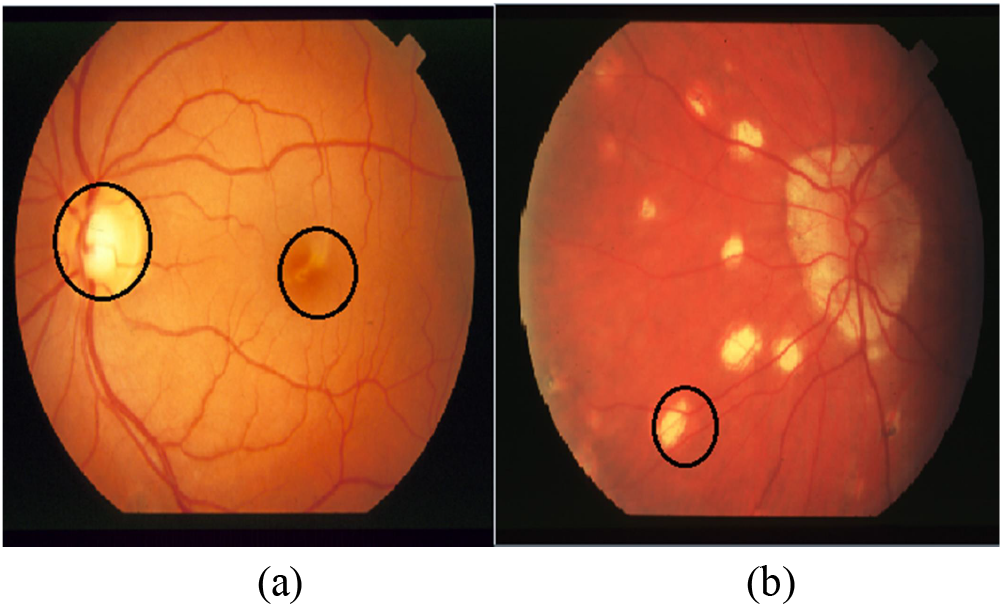
The figure 1(a) shows the OD and the macula. The figure 1(b) shows the exudate.

A normal optic disc is orange to pink in colour. A pale disc which varies in colour from orange colour to white is an indication of a disease condition. There is a similar bright region in the retina near the OD called the macula. Due to some eye diseases there are also other bright regions such as lesions, haemorrhage and exudates in the retina which makes the detection of OD a tedious process.

Several methods have been proposed to detect the OD. Some methods make use of the blood vessels to detect the OD by tracking the blood vessels [7– 11]. Some methods use the relative location between the macula and the OD that are often varying within a small range [12]. Some other method makes use of circular transformations to detect the OD [13–15]. In some methods a line operator is used which finds the OD based on the alignment of the line segments with minimum and maximum difference [6]. In some methods Hough transforms are used to detect circular regions [16–17]. But all these methods have complex algorithm to detect OD.

In our paper we have proposed an algorithm to detect the OD from other bright regions by using three characteristics of the OD. They are a) brightness b) circularity c) blood vessel. The OD is one of the brightest regions in the retina with highest intensity.

The OD is always circular in shape but exudates have irregular shape, their shape is not defined. In some cases we find that the exudates are circular and brighter than the OD. To differentiate the exudate from the OD, we generate a pattern similar to OD. We know that the convergence of blood vessels is maximum at the OD than in exudates. In the proposed algorithm we have undergone three levels of detection to locate the OD accurately.

## II. Proposed algorithm

In this section the proposed OD detection techniques is described. In particular, we divide the section into four subsections, which deals with finding maximum intensity, masking, normalised correlation, OD detection.

### A. Finding maximum intensity

Retinal images need to be pre-processed before finding the maximum intensity. Consider the retinal image I(x, y), from which we have to detect the OD. For that, firstly we have to convert the RGB image to grayscale image fig(2). Then determine the brightest region in the grayscale retinal image. In that we find regions with maximum intensity. The regions with maximum intensity determine the brightest points of the retinal image (I). The maximum intensity can be found as follows:

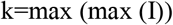

Where k contains the maximum intensities of the retinal image (I). Moreover, brightest pixel found using above formula can be present either in the OD of the retinal image or in the exudates. Further processes will help us to find the OD exactly in the retinal image.

**Figure2:**
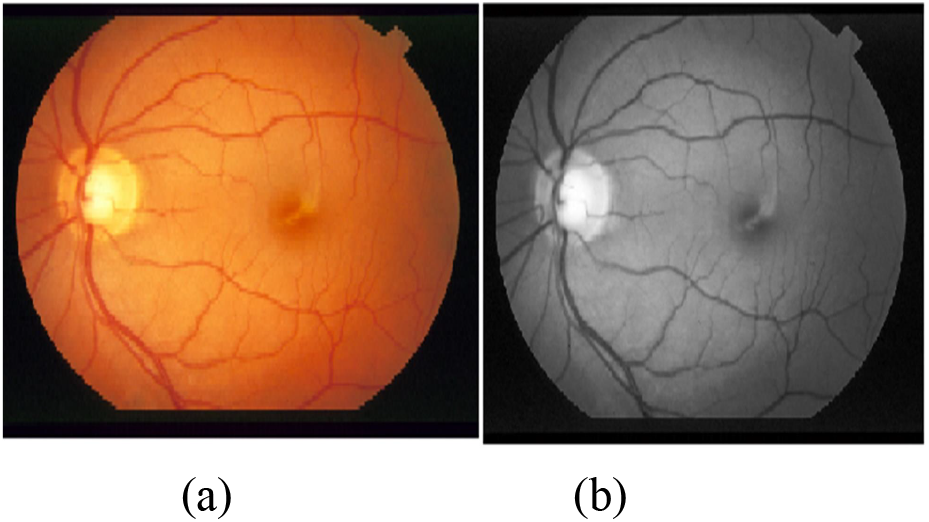
(a) is the retinal image and (b) is the grayscale image.

### B. Masking

In this technique the circular regions that have similar brightness structure as the OD are detected. The size of the input retinal image I is m × n. From that image, we take blocks of size j × j, where j=54, which is the standard size of circular OD [8]. We then find the blocks in which the length of k is maximum. The centre pixels of this block (27, 27) will be replaced by the M value calculated below.

To find the length of K in each block, we compare the pixel values of the particular block with the K value. The value of M can be estimated using the following formula:

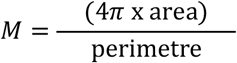

Where, Area is calculated as follows:

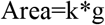

Where k is the maximum intensity and g represents the length of K of the block that of size 54 × 54. The value of M will be the minimum when the length of K is maximum. This indicates that the block is circular. The blocks where M value is zero, indicates that the block has less number of peak intensities or it does not have any peak intensity. So the value of g will be made minimum which in turn makes the M value zero. The perimeter to calculate M is taken as the block size 54, which is the standard OD size. Hence when the length of k is maximum it merely takes all the points of the block which represents circular shape.

This step is slide over the entire retinal image to generate the mask fig (3).The output mask image represents the places where the peak intensities are maximum and circular. Image may contain more than one area in case of the presence of exudates. Moreover the output image contains only zeros and ones. Also the output is having the same size of the input retinal image I(x, y).

**FIG 3:**
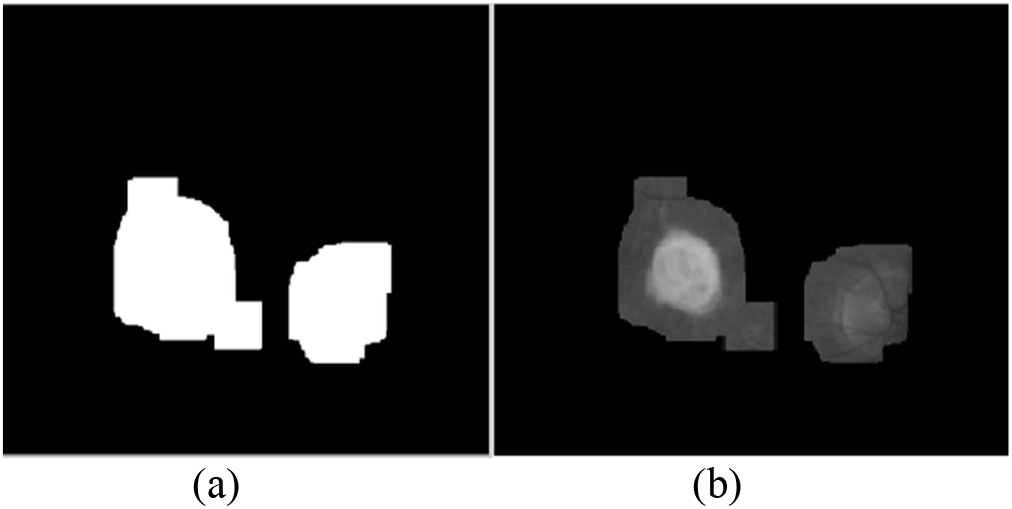
Fig 3(a) shows the mask obtained from M value and 3(b) shows the output of the mask on the input retinal image.

We apply the mask that we obtained in the earlier step, over the input retinal image in which the peak intensities are detected. The obtained image after replacing the mask image may contain OD and similar brighter regions like exudates. On applying the mask, exudates that are non-circular and small are not detected. This increases the chance of detecting the location of OD. But in some cases exudates that are circular are detected so we extend our algorithm to differentiate the OD and the exudate by the third characteristic feature of the OD blood vessels.

### C. Normalised Correlation

This approach is made to differentiate the OD exactly from the exudate. In most of the OD’s it is seen that the there is a central thick blood vessel and all other blood vessels branch from it. Moreover the convergence of blood vessel is maximum at the OD than in the exudate. We generate a pattern similar to the OD in which there are major blood vessels. Fig (4)

**Fig(4).**
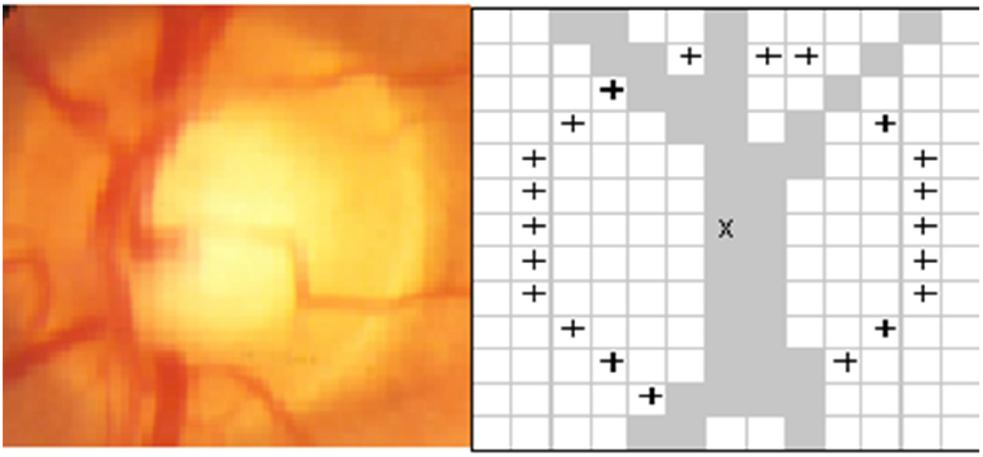
generated blood vessel pattern resembling OD

The generated pattern has the standard size of the OD (54 × 54). We now apply correlation between the image obtained from the mask and the pattern generated. We use Normalised 2D cross correlation on the image.

The cross correlation is given by,

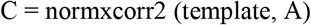

Where, A is the masked image.

C is the correlation coefficient matrix.

Template is the pattern generated fig (4). It is important that the size of the template must be smaller than the size of the mask image for better correlation. The correlation function is implemented using the equation as follows,

Correlation coefficient,

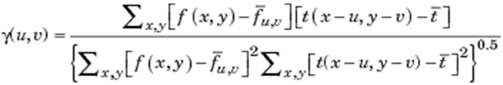

*f* is the image.

*t̄* is the mean of the template

*f̄*_*u,v*_ is the mean of *f*(*x,y*) in the region under the template.

### D. OD Detection

The values of the correlation coefficient ranges from −1 to 1. The correlation is the maximum at the points where the vessel pattern matches with the OD of the mask output. The correlation coefficient is the minimum at the exudates because it does not match with the pattern. The coordinates of the maximum correlation point is found. These coordinate points are replaced with a plus sign in the input retinal image. fig (5)

**Fig 5.**
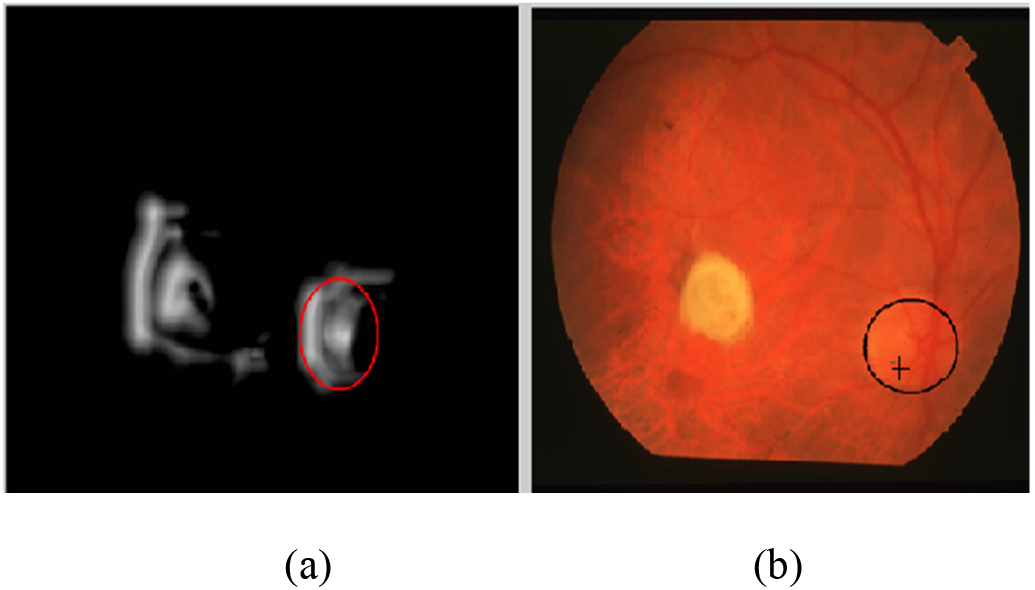
(a) shows the cross correlation output and (b) shows the exact location of the OD.

It is found that the plus sign detected the location of OD. From the above steps the location of OD is located exactly even when there are other bright regions in the image.

## III. Discussion

The proposed method exactly locates the OD from exudates and the macula. We experimented for proposed algorithm with the samples of DRIVE data set of various sizes. Out of 81 images of DRIVE dataset the algorithm located the OD exactly for 74 images. It failed for images where the OD is undefined. The accuracy of OD detection was 92%. The advantage of the proposed method is that it is not complex as in the previous methods of OD detection. The proposed algorithm is based on the features of the OD which makes the detection of OD location exactly.

The three important features of the OD that differs it from the other lesions present in the eye are the brightness, circularity and blood vessel pattern. To extract the brightness we converted the RGB colour image to grayscale in the green band, because grayscale images results in measuring the intensity of light at each pixel in a single band of electromagnetic spectrum. Since the OD is brighter we take the peak intensity points of the image. Since the OD is circular it makes clear that the OD location must have the maximum number of peak intensities. To separate the OD pixels from the image we go for masking the image. By the second level of OD detection we are able to detect the OD in 75%of the images. But the problem encountered was when there are retinal lesions and exudates as brighter as the OD and imperfectly circular, we found that the exudates were also detected along with the OD. In cases where the exudates are smaller in size than the OD only the OD is detected. Fig 6 shows the results of masking and the problems encountered in it.

**Fig 6:**
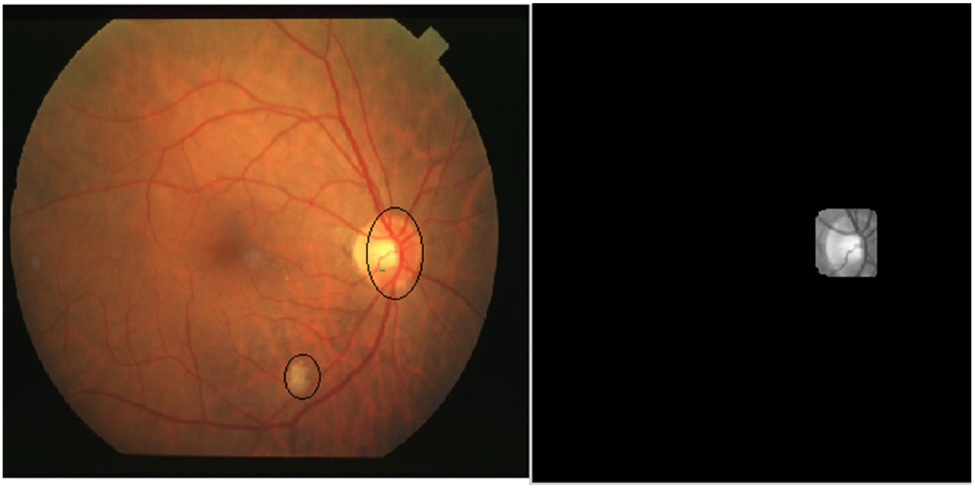
(a) the small exudates of the image are not masked.

**Fig 6:**
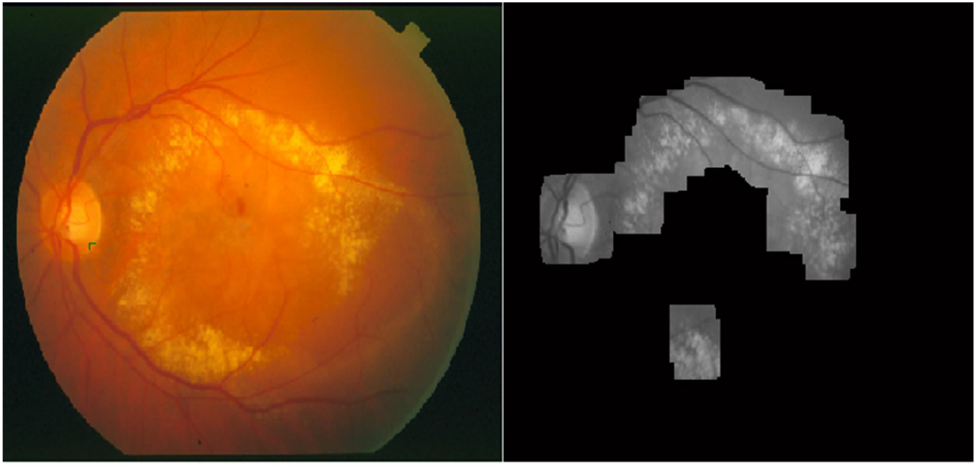
(b) the exudates that are bigger than the OD are masked.

To detect the OD from all these problems we extended our algorithm to the concept of correlation. We have seen that the major blood vessels converge at the OD than in the exudate. This is implemented using Normalized 2D Cross Correlation. The disadvantage of cross correlation is that, If the image energy Σ*f*^2^(*x,y*) varies with position the correlation between the feature and an exactly matching region in the image may be less than the correlation between the feature and a bright spot. The *correlation coefficient* overcomes these difficulties by normalizing the image and feature vectors to unit length, yielding a cosine-like correlation coefficient.

Normalized cross correlation is a mathematical computation that fulfils an essential role in image processing. Although it is well known that cross correlation can be efficiently implemented in the transform domain, the normalized form of cross correlation preferred for feature matching applications does not have a simple frequency domain. They are computed in the spatial domain. A template with blood vessels is generated for the correlation and the maximum correlation is obtained at the OD. The obtained results are shown.

## IV. RESULTS

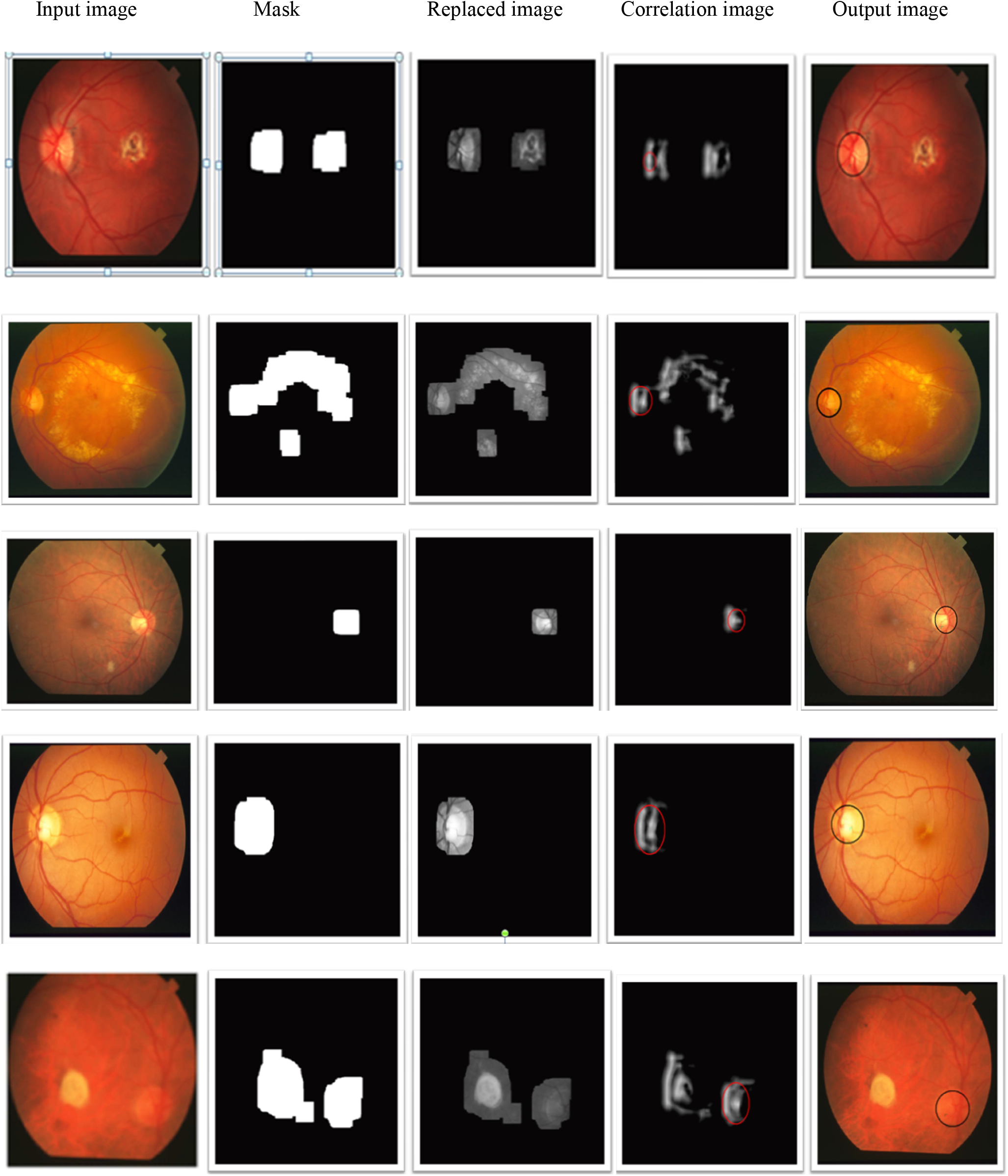

## V. CONCLUSION

This paper presents automatic OD detection from retinal images. With three levels of detection the OD is located. As in the previous detected methods this proposed algorithm is not so complex and the implementation is easy. Compared with the previous method the proposed technique does not require any transforms. The tolerant to different types of retinal lesions and imaging artefacts are proved through experiments done over drive database showed that the accuracy of 96% is obtained.

